# Effects of tumor mutation burden on the antigen presentation pathway

**DOI:** 10.1101/2021.04.14.439829

**Authors:** Enrique M. Garcia-Rivera, Jiho Park, Aakash Desai, Romain Boidot, Sandy Chevrier, Caroline Truntzer, François Ghiringhelli, Mitesh Borad, Aaron S. Mansfield

## Abstract

Tumor mutation burden (TMB) is used to select patients to receive immune checkpoint inhibitors (ICIs) but has mixed predictive capabilities. We hypothesized that inactivation of antigen presenting genes (APGs) that result from increased TMBs would result in inherent resistance to ICIs. We observed that somatic mutations in APGs were associated with increasing TMBs across 9,418 tumor samples of 33 different histological subtypes. In adenocarcinomas of the lung, *ITGAX* and *CD1B* were some of the most commonly mutated APGs. In 62 patients with non-small cell lung cancers treated with a PD-1 inhibitor in second or later lines of therapy, there was an association of increased TMB with mutations in APGs; however, mutations in one or more APGs were associated with improved progression-free survival. Contrary to our hypothesis, mutations in APGs were associated with improved progression-free survival with nivolumab, possibly due to the involvement of single alleles rather than complete loss.

## Introduction

Tumor mutation burden (TMB) has emerged as an important biomarker to select patients with cancer to treat with immune checkpoint inhibitors (ICI).^1^ With the application of next generation sequencing, TMB is frequently reported by commercial vendors and available for clinical interpretation. TMB has recently been accepted by the United States Food and Drug Administration as a biomarker to select patients to receive a PD-1 inhibitor based on the KEYNOTE-158 clinical trial.^1^ Mismatch repair deficiency, which is associated with high TMB, has also been approved to select patients to receive immunotherapy.^2^ Somatic mutations that are translated into mutant proteins have the potential to be presented by tumor cell major histocompatibility complex (MHC) class I proteins to CD8 T cells. Several retrospective and post hoc analyses have correlated TMB with response or survival in multiple tumor types.^3–5^

Acquired resistance to ICI has been reported to result from deleterious mutations in beta-2-microglobulin (*B2M*)^6^, which is critical for antigen presentation. Given the role that loss of antigen presentation has been implicated in acquired resistance to ICI, we hypothesized that increased TMB would more commonly involve antigen presenting genes (APG) and result in primary resistance to ICI. Accordingly, we sought to profile APG mutations across multiple tumor types and their association with TMB. We also sought to assess the outcomes of patients with mutant APG who were treated with ICI.

## Methods

Three sets of APGs were defined. The first set of genes was defined with the natural language processing tool *nferX Signals* by querying the term “antigen presentation” against a collection containing gene and proteins terms. Briefly, the tool we employed is based on word embedding models trained on biomedical literature in various corpora and supplemented with named entity recognition, synonyms handling, and temporal slicing. Our approach uses two key metrics – the Local Score (LS) and Global Score (GS) – to quantify the strength of literature association between two concepts across various corpora.^7^ Here we utilized the entirety of PubMed Abstracts and PMC Articles as our target corpus. There are 44 genes in this set coming from *nferX Signals* (**Supplemental Table 1**). The second set of genes was defined by one of the co-authors at Mayo Clinic, guided by prior publications.^8^ There are 45 genes in this set (**Supplemental Table 1**). The third set represented the union of the *nferX Signals* set and Mayo Clinic KOL set, consisting of 70 genes.

We analyzed 9,418 patient tumor samples in the The Cancer Genome Atlas (TCGA) dataset with somatic mutation and tumor mutational burden (TMB) data. The TCGA dataset comprises 33 different cancer histological subtypes. For each gene related to antigen presentation, we looked at the distribution of TMB values of patients with and without mutations in that gene on a pan-cancer basis, and within specific cancer types.

Descriptive statistics were used to summarize groups and outcomes. We quantified differences by computing the effect size (Cohen’s d) and p-values from a two-sided Mann-Whitney U test. Progression-free survival and overall survival were compared between patients with non-small cell lung cancer with and without mutations in APG treated with nivolumab^9^ using the log-rank test. This analysis was performed in R using the survival package.

## Results

To assess the associations of mutations in APGs with TMB, we first defined APGs. One APG set was defined using the *nferX Signals* application (nferX set = 44 genes), another set was defined by an author guided by a prior publication (Mayo set = 45 genes)^8^ and the union of the two sets (union = 70 genes) was also used.

When assessing 9,418 tumor samples from TCGA, mutations in one or more APGs had a large effect on the distribution of TMBs in the nferx set (Cohen’s d=0.731), the Mayo set (Cohen’s d=0.585) and the union of the two sets (Cohen’s d=0.710)(**Figures 1A-C**). Across all tumors, the most common mutations in APGs involved genes encoding human leukocyte antigens (**Table 1**). Mutations in *TAP2* and *HLA-DRB9* had the greatest effects on increased TMB (Cohen’s d= 2.46 and 2.86, respectively; **Figure 1D**). When assessing individual mutations, there was a large increase in the distribution of TMB values of patients with the *TAP2* delC mutation (chr6:32838010) which was present in 0.15% of cases (Cohen’s d=2.574). There were also large increases in the distributions of TMB values of patients with the *CANX* delT mutation (chr5:179722918) which was present in 0.19% of cases (Cohen’s d=2.233), and the *B2M* delCT mutation (chr15:44711582) which was present in 0.31% of cases (Cohen’s d=1.724).

**Table 1:** List of most frequently mutated antigen presenting genes across all tumors and their effects on tumor mutation burden

| Gene | Position | Base change | Frequency | Cohen's d |
| --- | --- | --- | --- | --- |
| *HLA-A* | chr6:29942911 | A>G | 2.34% | 0.05 |
| *HLA-DRB1* | chr6:32581806 | A>G | 2.33% | 0.01 |
| *HLA-A* | chr6:29942916 | A>G | 1.79% | -0.01 |
| *HLA-DRB5* | chr6:32530123 | T>C | 1.72% | 0.01 |
| *HLA-DRB1* | chr6:32581805 | C>A | 1.64% | -0.01 |
| *HLA-DRB1* | chr6:32580855 | A>T | 1.55% | -0.06 |
| *HLA-DRB1* | chr6:32580767 | A>C | 1.47% | 0.02 |
| *HLA-C* | chr6:31271132 | C>T | 1.45% | 0.06 |
| *HLA-DRB5* | chr6:32530128 | C>T | 1.39% | 0.32 |
| *HLA-DQB1* | chr6:31271133 | A>C | 1.31% | 0.12 |
| *HLA-C* | chr6:31271133 | A>C | 1.31% | 0.20 |
| *HLA-DRB1* | chr6:32581763 | G>A | 1.26% | 0.32 |
| *HLA-C* | chr6:31271153 | T>G | 1.05% | 0.12 |

**Figure 1:**
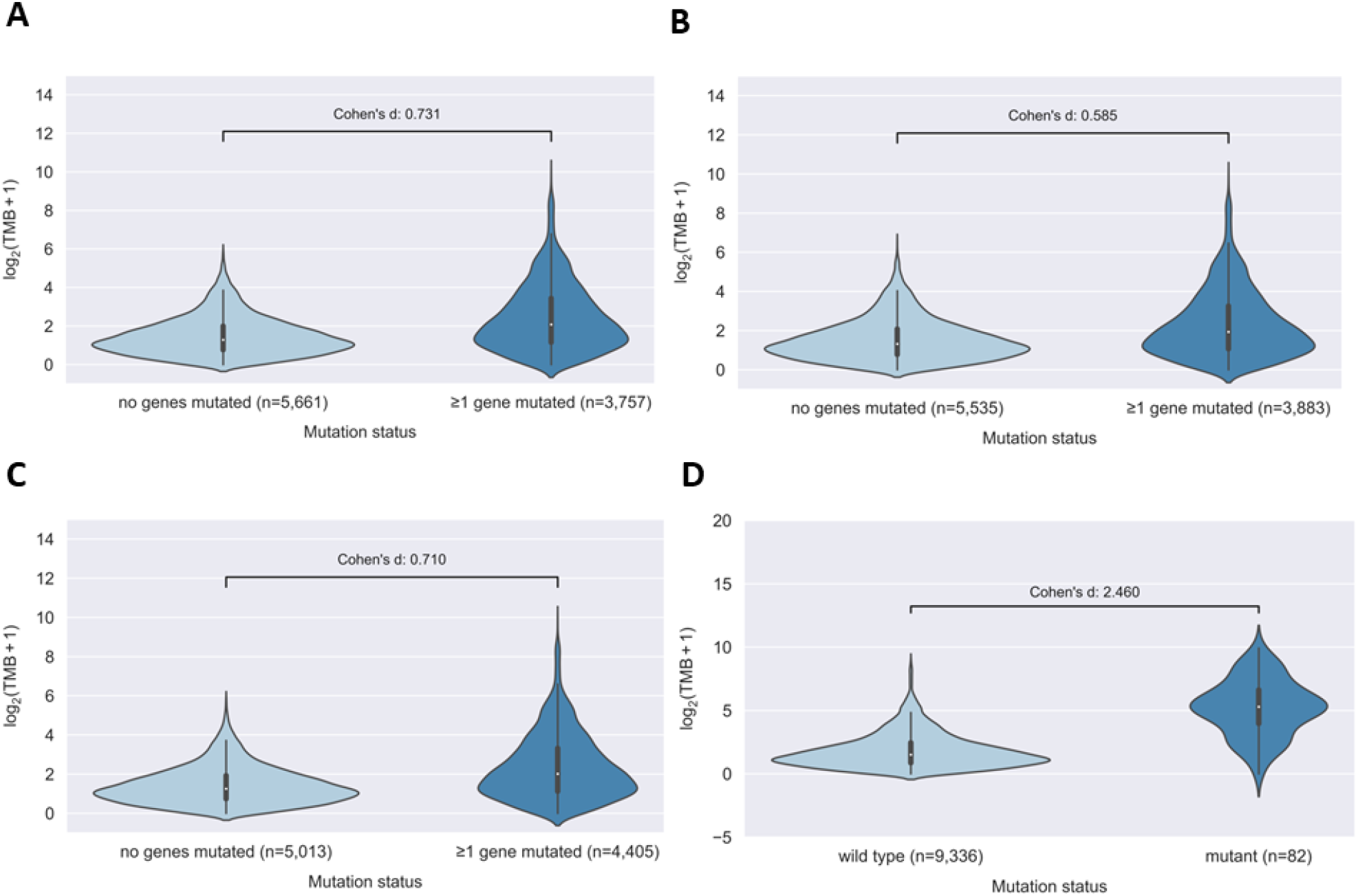
The distribution of tumor mutation burdens (TMB) are shown by whether mutations in antigen presenting genes (APG) are present or not across the TCGA dataset based on the nferx (A), Mayo (B) and the union of both (C) APG lists. TMBs are also shown by whether mutations in *TAP2*, an antigen peptide transporter, was present or not (D).

There was significant variation in the effects of mutations in APGs on the distributions of TMBs across specific cancer types when looking at the union of both gene sets (**Figure 2A**), with the largest effect sizes seen in uterine corpus endometrial carcinomas and diffuse large B cell lymphomas. Meanwhile, negative effect sizes were seen in tenosynovial giant cell tumors and renal papillary cell carcinomas. In adenocarcinomas of the lung, *ITGAX*, a gene encoding an integrin, was one of the most frequently mutated APGs with a large effect on the distribution of TMBs (**Figure 2B, Table 2**). *CIITA*, a transcriptional coactivator, had one of the largest effect sizes on the distribution of TMBs in adenocarcinomas of the colon (**Figure 2C, Table 3**). LY75, an endocytic receptor, had one of the largest effect sizes on the distribution of TMBs in melanoma (**Figure 2D, Table 4**).

**Table 2:** Top ten genes with the greatest effect size on TMB in lung adenocarcinomas

| Gene | Mutation frequency | Patients with mutation | Cohen's d | Neg log U-test p-value |
| --- | --- | --- | --- | --- |
| *CD80* | 0.39% | 2 | 1.75 | 1.58 |
| *PSME2* | 0.39% | 2 | 1.70 | 1.56 |
| *CD83* | 0.39% | 2 | 1.61 | 1.48 |
| *HLA-E* | 0.39% | 2 | 1.54 | 1.35 |
| *PDIA3* | 0.98% | 5 | 1.51 | 2.60 |
| *HLA-DMB* | 0.39% | 2 | 1.41 | 1.10 |
| *B2M* | 1.77% | 9 | 1.19 | 3.11 |
| *ITGAX* | 8.66% | 44 | 1.13 | 10.58 |
| *IL2* | 0.98% | 5 | 1.12 | 1.85 |
| *CD1B* | 2.56% | 13 | 1.10 | 3.96 |

**Table 3:** Top ten genes with the greatest effect size on TMB in colon adenocarcinoma

| Gene | Mutation frequency | Patients with mutation | Cohen's d | Neg log t-test p-value | Neg log U-test p-value |
| --- | --- | --- | --- | --- | --- |
| *HLA-DOB* | 0.55% | 2 | 3.21 | 1.17 | 1.79 |
| *CD80* | 0.83% | 3 | 3.08 | 1.56 | 2.36 |
| *CD86* | 0.83% | 3 | 2.48 | 0.70 | 1.68 |
| *PDIA3* | 3.04% | 11 | 2.48 | 4.59 | 5.84 |
| *CD1C* | 1.38% | 5 | 2.43 | 1.55 | 2.69 |
| *CTSS* | 1.38% | 5 | 2.39 | 1.48 | 2.40 |
| *IFI30* | 0.83% | 3 | 2.39 | 0.64 | 1.12 |
| *IRF1* | 1.93% | 7 | 2.20 | 2.17 | 3.14 |
| *CANX* | 2.21% | 8 | 2.16 | 2.90 | 3.66 |
| *CIITA* | 6.08% | 22 | 2.16 | 8.00 | 8.36 |

**Table 4:** Top ten genes with the greatest effect size on TMB in cutaneous melanoma

| Gene | Mutation frequency | Patients with mutation | Cohen's d | Neg log U-test p-value |
| --- | --- | --- | --- | --- |
| *TAPBP* | 0.43% | 2 | 2.64 | 1.73 |
| *FCGRT* | 1.07% | 5 | 2.10 | 3.06 |
| *CSF2* | 0.85% | 4 | 2.07 | 2.12 |
| *IL2* | 0.85% | 4 | 1.61 | 1.33 |
| *PSME2* | 1.71% | 8 | 1.55 | 3.21 |
| *CD74* | 1.49% | 7 | 1.54 | 2.44 |
| *LY75* | 8.96% | 42 | 1.48 | 14.37 |
| *HLA-G* | 1.49% | 7 | 1.46 | 2.92 |
| *CANX* | 0.85% | 4 | 1.46 | 1.52 |
| *PSMB8* | 1.28% | 6 | 1.43 | 2.83 |

**Figure 2:**
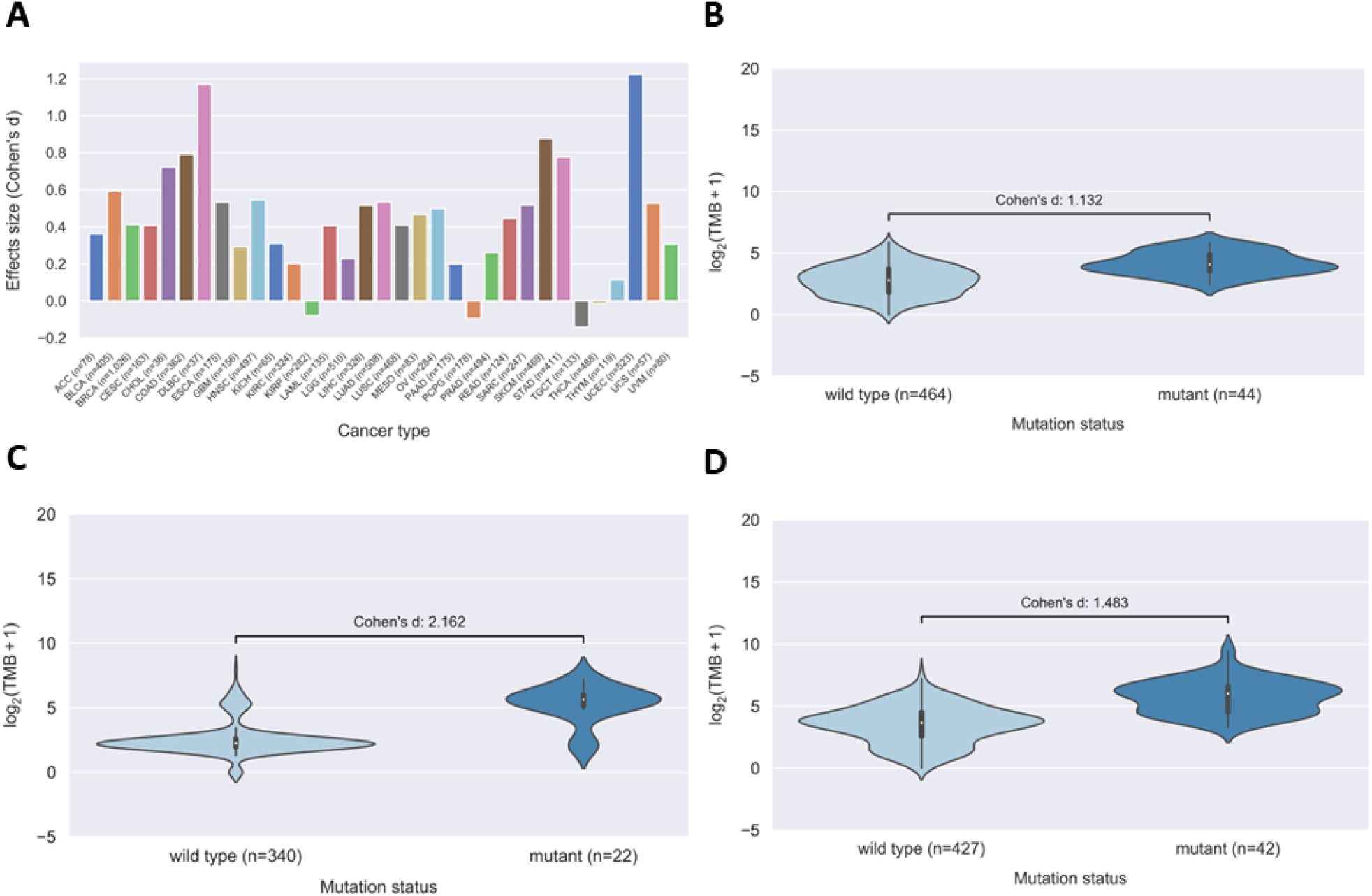
The effect sizes (Cohen’s d) of the associations of TMB with mutations in APGs (from the union of both gene sets) is shown across the tumor types (A). When looking at the effect sizes of mutations in specific APGs with TMB, mutations in ITGAX (B), CIITA (C) and LY75 (D) had the greatest effect sizes in adenocarincoma of the lung, colorectal cancer and melanoma respectively.

To assess whether mutations in APGs affected survival in non-small cell lung cancer, we analyzed 62 patients who received the PD-1 inhibitor nivolumab as a second or later line of therapy and had paired somatic and germline sequencing data available.^9^ Of these patients, 72.6% were male and the median age was 66 years. The median progression free survival was 2.3 months and the median overall survival was 12.2 months. There were thirteen tumors with KRAS mutations, four with *EGFR* mutations and none with *ALK* rearrangements. Thirty-five patients (56%) had one or more mutations in APGs, whereas 27 patients (44%) had no mutations in APGs. Most mutations in APGs were not bi-allelic. Prior analysis of this dataset demonstrated improvements in progression free survival in patients whose tumors had high tumor mutations burdens. We analyzed survival based on the presence or absence of mutations in APGs. There was no difference in overall survival between patients with one or more mutations in APGs compared to those with wild type APGs (median OS: 21.6 v 13.3 months, HR 0.78, 95%CI 0.39-1.57, p=0.49; Figure 3A). In contrast, progression-free survival was superior in patients with one or more mutations in APGs compared to those with wild type APGs (median PFS: 3.7 v 1.8 months; HR=0.54, 95%CI 0.30-0.97, p=0.04; **Figure 3B**).

**Figure 3:**
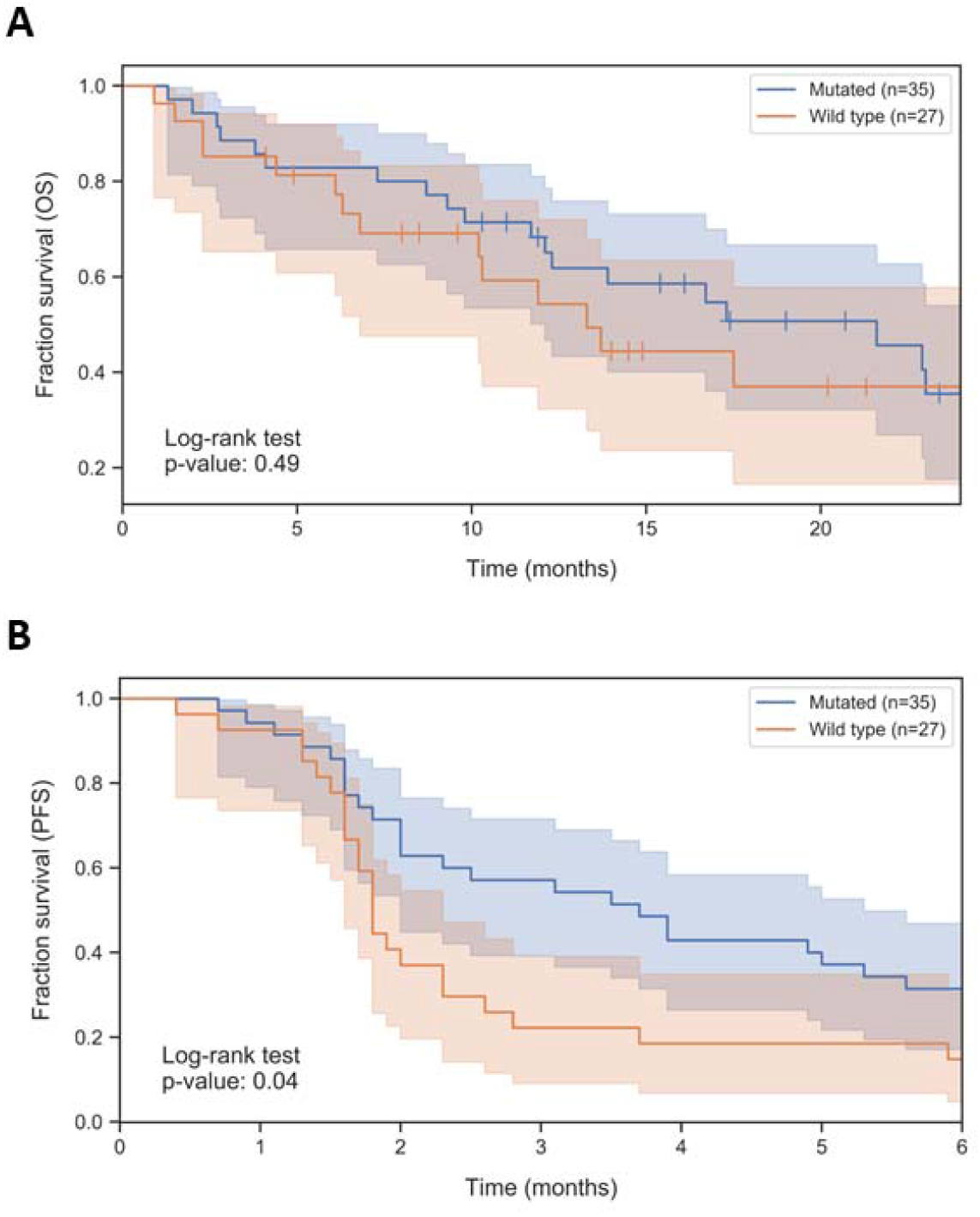
In a cohort of 62 patients who received the PD-1 inhibitor nivolumab as second or later lines of therapy, the overall survival (A) and progression-free survival (B) are shown between patients with and without mutations in APGs.

## Discussion

In our pan-cancer analysis, the distributions of TMB were significantly higher in specimens with mutations in one or more in APGs. Among all tumor types, this effect was largest in uterine corpus endometrial carcinomas and diffuse large B cell lymphomas. Overall, mutations in genes that encode human leukocyte antigens were the most frequently involved APGs; however, mutations in *TAP2* and *HLA-DRB9* were associated with the largest increases in TMB. Contrary to our hypothesis, there was no survival benefit in patients with non-small cell lung cancer and mutant APGs treated with the PD-1 inhibitor nivolumab.

Similar to our findings that mutations in *TAP2* had one of the greatest effects on TMB across all tumors, others have reported that mutations in bare lymphocyte syndrome genes (which include *TAP2*) are associated with high TMB and neoantigen loads.^10^ Other genes associated with bare lymphocyte syndrome including *CIITA* and TAPBP also had large effects on TMB in our analysis. Primary or acquired resistance to immunotherapy may occur due to somatic loss of antigen presentation, similar to severe immunodeficiency caused by germline loss of antigen presentation.

Beta-2-microglobulin is necessary for peptide-major histocompatibility complex class I formation at the cell surface. Mutations in *B2M* have been associated with acquired resistance to PD-1 inhibitors.^6^ Others have replicated these findings, demonstrating that mutations in *B2M* are associated with acquired resistance to anti-CTLA-4 and anti-PD-1 therapies in patients with melanoma, and additionally suggested that primary resistance to treatment can also result from mutations in *B2M*.^11^ Previous studies have shown that cells exposed to strong T cell-mediated selective pressure may develop preferential molecular defects in B2M gene. For example, the *B2M* delCT mutation is preferentially selected in melanoma cells after T cell-based immunotherapy.^12^ We also found that mutations in *B2M* were associated with higher TMB.

Given the associations with mutations in APGs and acquired resistance to immunotherapies, we hypothesized that primary resistance could also be related to mutations in APGs. Although we saw that mutations in APGs were associated with large increases in TMB, our analysis of a cohort of patients with NSCLC identified a benefit in progression free survival when APGs were mutated, contrary to our initial hypothesis.

Upon further inspection, most of the mutations in this cohort only involved one allele, suggesting that the adequate antigen presentation was possible in these cases with a preserved wild type allele. Previous reports suggest bi-allelic disruption of APGs minimizes immunogenicity in human embryonic stem cells.^13^ Furthermore, given the association we identified with mutations in APGs and TMB, our analysis was aligned with the original report of this cohort demonstrating improved progression free survival in the patients with high TMB.

Retrospective and post-hoc analyses have suggested that tumor mutation burden is a predictor of response to or survival with immunotherapy.^3–5^ There are significant discrepancies in the scoring of TMB in some tumor types, such as renal cell carcinoma where insertions and deletions have a greater effect on TMB than single nucleotide variants.^14^ Additionally, the combination of ipilimumab and nivolumab was recently approved for mesothelioma based on improvements in survival even though these tumors typically have the lowest mutation burdens of any carcinogen-related malignancy.^15, 16^ Recently, chromosomal rearrangements were reported to have neo-antigenic potential in mesothelioma^17^ and head and neck cancers.^18^ These studies imply that insertions, deletions and rearrangements could improve upon TMB scores that only include single nucleotide variants. It has also been suggested that HLA-correction^19, 20^ and germline subtraction^21^ may further improve TMB scoring. Given the lack of a standardized approach to calculate TMB, there are calls to harmonize and standardize the methodology.^22–24^

In conclusion, we found that mutations in APGs are associated with increases in tumor mutation burdens. Although APG mutations were not found to affect the response to immunotherapy, bi-allelic loss of genes critical to antigen presentation such as *B2M* may result in primary or acquired resistance to immunotherapies. As our understanding of TMB and efforts to standardize TMB scoring continue to grow, attention to loss of APGs may further inform these efforts.

## Data Availability

Data from TCGA may be accessed per instructions here: https://gdc.cancer.gov/gdc-tcga-data-access-matrix-users. The survival and genomic data from the Dijon cohort are available by request from the authors of the original work.^9^

## Acknowledgements

The authors would like to thank BobbiAnn Jebens for her editorial assistance. This work was supported in part by nference, Mayo Clinic’s Center for Individualized Medicine, NCI R21 CA251923 and a Mark Foundation ASPIRE award.

## Author Contributions

Conceived and designed the analysis: EG, ASM

Collected the data: EG, JP, CT, FG, MB, ASM

Contributed data or analysis tools: CT, FG

Performed the analysis: EG, JP, CT, FG, MB, ASM

Wrote the paper: EG, JP, CT, FG, MB, ASM

---

**Supplemental table 1:** Antigen presentation gene lists

| nferx | Mayo KOL |
| --- | --- |
| *APCS, CALR, CD1A, CD1B, CD1C, CD1D, CD1E, CD209, CD40, CD74, CD80, CD83, CD86, CIITA, CPQ, CSF2, CTSS, ERAP1, ERAP2, FCER2, FCGRT, HLA-A, HLA-B, HLA-C, HLA-DMA, HLA-DMB, HLA-DPA1, HLA-DPB1, HLA-DQA1, HLA-DQB1, HLA-DRA, HLA-DRB1, HLA-E, IFI30, IL10, IL2, ITGAX, LY75, PSMA1, PSMB8, PSMB9, TAP1, TAP2, TAPBP* | *B2M, CALR, CANX, CIITA, ERAP1, ERAP2, HLA-A, HLA-B, HLA-C, HLA-DMA, HLA-DMB, HLA-DOA, HLA-DOB, HLA-DPA1, HLA-DPA2, HLA-DPB1, HLADPB2, HLA-DQA1, HLA-DQA2, HLA-DQB1, HLA-DQB2, HLA-DRA, HLA-DRB1, HLA-DRB5, HLA-DRB9, HLA-E, HLA-F, HLA-F-AS1, HLA-G, HLA-J, HLA-K, HLA-L, HLA-P, HLA-W, HSPBP1, IRF1, NLRC5, PDIA3, PSMA7, PSME1, PSME2, PSME3, TAP1, TAP2, and TAPBP* |

